# The Nuclear And Mitochondrial Genomes Of The Facultatively Eusocial Orchid Bee *Euglossa dilemma*

**DOI:** 10.1101/123687

**Authors:** Philipp Brand, Nicholas Saleh, Hailin Pan, Cai Li, Karen M. Kapheim, Santiago R. Ramírez

## Abstract

Bees provide indispensable pollination services to both agricultural crops and wild plant populations, and several species of bees have become important models for the study of learning and memory, plant-insect interactions and social behavior. Orchid bees (Apidae: Euglossini) are especially important to the fields of pollination ecology, evolution, and species conservation. Here we report the nuclear and mitochondrial genome sequences of the orchid bee *Euglossa dilemma* Bembé & Eltz. *Euglossa dilemma* was selected because it is widely distributed, highly abundant, and it was recently naturalized in the southeastern United States. We provide a high-quality assembly of the 3.3 giga-base genome, and an official gene set of 15,904 gene annotations. We find high conservation of gene synteny with the honey bee throughout 80 million years of divergence time. This genomic resource represents the first draft genome of the orchid bee genus *Euglossa,* and the first draft orchid bee mitochondrial genome, thus representing a valuable resource to the research community.

## Introduction

Bees (Apoidea) are important models for the study of learning and memory (Menzel and Muller 1996), plant-insect interactions (Doetterl and Vereecken 2010) and the evolution of social behavior (Nowak *et al*. 2010; Woodard *et al. 2011*; Kapheim *et al*. 2015). Among the >20,000 bee species worldwide, lineages have evolved varied degrees of specialization on floral resources such as pollen, resins, and oils (Wcislo and Cane 2003; Michener 2007; Litman *et al*. 2011). These relationships are wide-ranging and have substantial impact on the health and function of natural and agricultural systems (Klein *et al*. 2007). Furthermore, several transitions from an ancestral solitary to a derived eusocial behavior have occurred within bees (Danforth 2002; Cardinal and Danforth 2011; Branstetter *et al*. 2017). Thus, bees provide unique opportunities to investigate the genetic underpinnings of multiple major ecological and evolutionary transitions. The repeated evolution of different behavioral phenotypes in bees, including foraging and social behavior, provides a natural experiment that allows for the determination of general as well as species-specific molecular genomic changes underlying phenotypic transitions. In order to capitalize on this potential, whole-genome sequences of a divergent array of bee species with different life histories are needed (Kapheim *et al*. 2015).

Orchid bees (Apidae; Euglossini) are among the most important pollinators of angiosperms in the neotropical region (Ramírez *et al*. 2002). While female orchid bees collect nectar, pollen and resin for nest construction and brood-cell provisioning, male bees collect perfume compounds from floral and non-floral sources (Vogel 1966; Whitten *et al*. 1993; Eltz *et al*. 1999; Roubik and Hanson 2004). These volatile compounds are used to concoct a species-specific perfume blend that is subsequently used during courtship display, presumably to attract conspecific females. This unique male scent-collecting behavior has recently been examined in a broad array of molecular ecological and evolutionary studies, focusing on phenotypic evolution, chemical communication, plant-insect mutualisms, and speciation (Eltz *et al.* 2008; 2011; Ramírez *et al.* 2011; Brand *et al.* 2015; Weber *et al.* 2016).

While most of the approximately 220 species of orchid bees appear to be solitary, several species have transitioned to living in coordinated social groups (Garófalo 1985; Pech *et al.* 2008; Augusto and Garófalo 2009). Female *Euglossa dilemma* individuals, for example, can either form solitary nests and provision their own brood cells, or live in small groups where daughters remain in their natal nest and help their mother rear her offspring, instead of dispersing to found their own nest. The social *E. dilemma* nests (similar to the closely related *E. viridissima;* Pech *et al.* 2008) exhibit true division of labor, with subordinate daughters foraging for resources and the reproductively dominant mothers remaining in the nest and laying eggs (Saleh & Ramírez pers. obs.). This facultative eusocial behavior represents an early stage in social evolution and makes *E. dilemma* well suited for studying the genetic mechanisms underlying the transition from solitary to eusocial behavior. While other facultative eusocial species evolved throughout the bee lineage, orchid bees have a unique taxonomic position (Cardinal and Danforth 2011). Orchid bees are part of the corbiculate bees, together with the honey bees, bumble bees and stingless bees, three obligately eusocial bee lineages (Figure 1a). As the sister group to all other corbiculate bee lineages (Romiguier *et al.* 2016; Branstetter *et al.* 2017; Peters *et al.* 2017), orchid bees may provide key insights into the early stages of eusociality and the possible evolutionary trajectories that led to the obligate eusocial behavior exhibited by honey bees.

**Figure 1.**
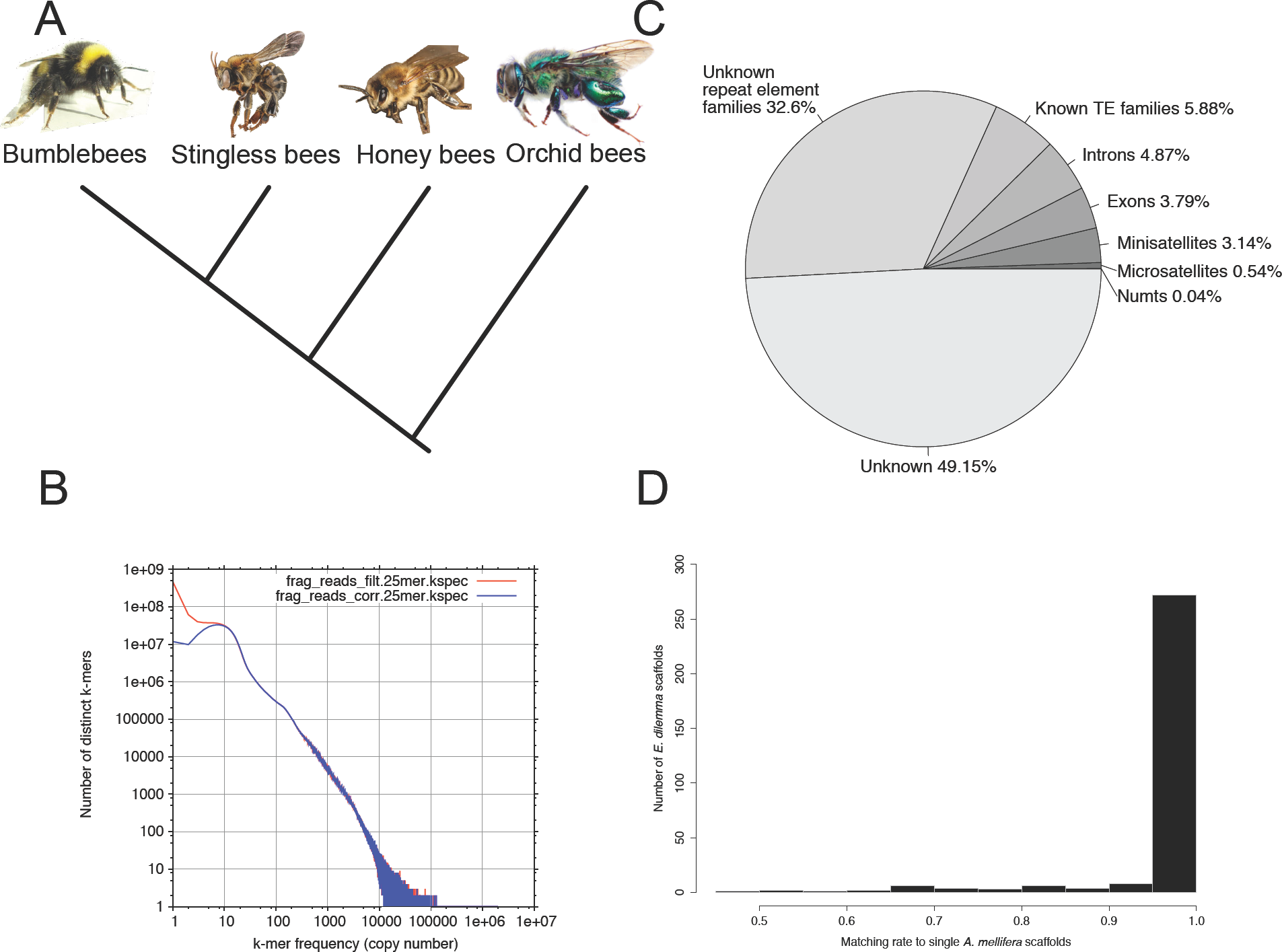
Genomic features. **(A)** Phylogeny of the four corbiculate bee tribes with orchid bees as sistergroup to honey bees, stingless bees, and bumblebees (Romiguer et al. 2015). **(B)** K-mer distribution spectrum (k=25) of genomic sequence reads. The positively skewed spectrum reveals a high abundance of a few k-mers, leading to an estimate of 87.7% repetitiveness of the *E. dilemma* genome. Red shows the kmer spectrum before, and blue after error correction. **(C)** Genomic element density including genic and non-genic features as a fraction of the overall genome assembly length excluding stretches of N. Over 49.15% of the assembly could not be annotated with the selected methods. **(D)** Synteny between the *E. dilemma* and the honey bee (*Apis mellifera*) genome. In an analysis including *E. dilemma* scaffolds of ≥100kb length, 83% showed ≥95% synteny to a single honey bee scaffold. Photographs in **(A)** are reproduced from Wikimedia under the CC BY-SA 3.0 license.

Here we present the draft genome of the orchid bee species *Euglossa dilemma*. Using a combined paired-end and mate-pair library sequencing approach, we assembled 18% of the predicted 3.3Gb genome, and annotated a high-quality gene set including 15,904 genes. In addition, we reconstructed three quarters of the mitochondrial genome with the help of transcriptome data, representing the first orchid bee mitogenome. These genomic resources will facilitate the genetic study of outstanding ecological and evolutionary questions, such as the evolution of resource preferences and the evolution of eusociality. Moreover, it provides an important genomic resource for a group of crucial neotropical pollinators, that are of specific concern for conservation biologists (Zimmermann *et al.* 2011; Suni and Brosi 2012; Suni 2016; Soro *et al.* 2016).

## Materials and Methods

### Genome sequencing and assembly

#### Nuclear Genome

Sequencing of the *E. dilemma* genome was based on six haploid male individuals collected at Fern Forest Nature Center in Broward County, FL(26°13’46.3’’N, 80°11’08.9’’W) in February 2011. This population was chosen based on its low nucleotide diversity resulting from a bottleneck during a single introduction to Southern Florida about 15 years ago (Skov and Wiley 2005; Pemberton and Wheeler 2006; Zimmermann *et al.* 2011). DNA was extracted from each bee independently and used for the construction of four paired-end (two 170bp and 500bp libraries, respectively) and four mate-pair (two 2kb and 5kb libraries, respectively) sequencing libraries. Next, the paired-end libraries were sequenced in 90 cycles and the mate-pair libraries for 49 cycles on an Illumina HiSeq2000. The resulting sequence data was run through fastuniq v1.1 (Xu *et al.* 2012) to remove PCR duplicates and quality trimmed using trim_galore v0.3.7 (Babraham Bioinformatics). Subsequently, reads were used for *de novo* assembly with ALLPATHS-LG v51750 (Gnerre *et al.* 2011) and Soap-denovo2 (Luo *et al.* 2012) with varying settings. Gaps within scaffolds were closed using GapCloser v1.12 (Luo *et al.* 2012) for each assembly. ALLPATHS-LG with default settings resulted in the highest-quality assembly, based on assessments of annotation completeness (see below). This assembly (*E. dilemma* genome assembly v1.0) was used for all subsequent analyses. All other assemblies were excluded from analysis, but are available upon request.

The pre-processed reads were used for k-mer based genome size estimates. We used ALLPATHS-LG to produce and analyze the k-mer frequency spectrum (k=25). Genome size was estimated on the basis of the consecutive length of all reads divided by the overall sequencing depth as (*N* × (*L* – *K* + 1) – *B*)/*D* = *G*, where *N* is the total number of reads, *L* is the single-read length, *K* is the k-mer length, *B* is the total count of low-frequency (frequency ≤3) k-mers that are most likely due to sequencing errors, *D* is the k-mer depth estimated from the k-mer frequency spectrum, and *G* is the genome size. In addition, we used the ALLPATHS-LG k-mer frequency spectrum to predict the repetitive fraction of the genome.

The quality of the genome assembly was assessed using standard N statistics, and assembly completeness as measured by the CEGMA v2.5 (Parra *et al.* 2007) and BUSCO v1.1 (Simão *et al.* 2015) pipelines. CEGMA was run in default mode, whereas BUSCO was run with the arthropoda_odb9 OrthoDB database (Zdobnov *et al.* 2017) in genome mode.

We estimated the mean per-base genome coverage on the basis of the pre-processed reads and the estimated genome size as 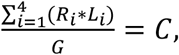 where *R* is the number of reads and *L* the mean read length of sequence library *i*, *G* is the estimated genome size and *C* the resulting per-base coverage.

#### Mitogenome

Initial attempts to reconstruct the mitochondrial genome from our whole-genome shotgun sequencing reads including read subsampling and exclusion of rare variants were only partially successful, due to high sequence variability of sequencing reads with similarity to mitochondrial loci (data not shown). In addition, we have observed that the amplification of mitochondrial DNA in standard polymerase chain reactions (PCR) leads to a high level of polymorphic sites in *E. dilemma* and other orchid bees (Brand & Ramírez pers. obs.). Together, this suggests the presence of nuclear copies of the mitochondrial genome (NUMTs) that interfere with the assembly process and PCR amplification. Mitochondrial genes are expressed in almost every tissue in eukaryotes. We used this feature to reconstruct the mitochondrial genome as far as possible from RNA-Seq data. Therefore, we used available *E. dilemma* transcriptome assemblies in order to reconstruct the mitochondrial genome from cDNA (Brand *et al.* 2015). In order to find expressed mitochondrial genes represented in the transcriptome assembly of Brand et al. 2015, we used blastx with the honey bee mitochondrial genome as query (Crozier and Crozier 1993) and an E-value cutoff of 10E-12 (Altschul *et al.* 1990; Camacho *et al.* 2009). The contigs and scaffolds that were detected with this approach were annotated following (Dietz *et al.* 2016). Briefly, we performed tblastn and tblastx searches with protein coding genes and rRNA genes of the honey bee mitochondrial genome, respectively. All hits were used for manual gene annotation using Geneious v8.0.5 (Biomatters Ltd. 2012). Since the recovered mitochondrial mRNA scaffolds contained more than one gene, we searched and annotated intergenic tRNAs using ARWEN 1.2.3 (Laslett and Canbäck 2008) and tRNAscan-SE 1.21 (Lowe and Eddy 1997).

### Genome annotation

#### Gene annotation

Genes were annotated based on sequence homology and *de novo* gene predictions. The homology approach was based on the recently updated high-quality official gene set of the honey bee (OGS v3.2; Elsik *et al.* 2014). All honey bee original gene set (OGS) proteins were used in initial tblastn searches against all *E. dilemma* scaffolds with an E-value cutoff of 10E-4. All honey bee protein sequences with a blast hit to the *E. dilemma* genome assembly covering at least 50% of the protein were selected for homology based annotation. The resulting set of honey bee proteins was used as input to exonerate v2.42.1 (Slater and Birney 2005) in order to annotate homologous open reading frames (ORFs) through accurate exonintron boundary prediction for each scaffold. Exonerate was run with default settings and the minimum fraction of the possible optimal similarity per protein query set to 35%. In a second round, genes not annotated under the previous settings were rerun with minimum similarity set to 15%. In the case of overlapping annotations on the same strand (*i.e.* identical ORF orientation) resulting from honey bee queries with high similarity, we discarded all but one annotation with the best exonerate score (based on completeness and similarity). This approach proved feasible due to the relatedness of *E. dilemma* and the honey bee, as well as the high quality of the well curated honey bee OGS. For *de novo* gene prediction we used Augustus (Stanke *et al.* 2008) and SNAP (Korf 2004) trained on the honey bee, with the *E. dilemma* genome masked for repetitive regions (See below) as input. Only genes predicted by both programs were taken into account. Gene predictions with ≥85% sequence similarity to each other were discarded, to prevent the inclusion of putative unmasked transposable element derived genes in the official gene set. *De novo* predictions were added to the *E. dilemma* OGS if not annotated by the homology-based approach.

#### Repetitive element annotation

Repetitive elements including tandem repeats, nuclear copies of the mitochondrial genome (NUMTs), and transposable elements (TEs) were annotated using multiple methods.

#### Tandem repeats

We searched for micro- and mini-satellites (1–6 bp and 7–1000 bp motif length, respectively) in all scaffolds using Phobos 3.3.12 (Mayer 2010). We performed two independent runs for each class of tandem repeats with Phobos parameter settings following Leese et al. 2012 (gap score and mismatch score set to -4 and a minimum repeat score of 12; Leese *et al.* 2012).

#### NUMTs

We annotated NUMTs using blastn runs with the partial mitochondrial genome (see above) as query and an E-value cutoff of 10E-4 as used in NUMT analyses of other insect genomes (Pamilo *et al.* 2007). This approach allowed us to find NUMTs with medium to high similarity to the actual transcriptome-based mitochondrial genome.

#### TEs

In order to annotate TEs, we first ran RepeatModeler for *de novo* repeat element annotation and classification based on the genome annotation followed by RepeatMasker in order to detect the total fraction of repetitive elements present in the genome (Smit et al. 2016). In addition we used the k-mer based *de novo* repeat assembler REPdenovo (Chu *et al.* 2016) to identify highly repetitive genomic elements based on the generated short sequence reads. These two methods independently assess repetitive sequence content in genomic data using different approaches thus allowing for a more robust estimation of repetitive genome content. The incorporation of multiple *de novo* repeat detection pipelines was necessary due to the lack of a bee-specific repeat database. RepeatModeler v1.0.8 was run with default settings using the ncbi blast algorithm (Altschul *et al.* 1990) for repeat element detection. We used the resulting *de novo* repeat element annotations as a database for Repeatmasker v4.0.5 with Crossmatch v0.990329 as the search engine. We ran the analysis in sensitive mode in order to identify the total fraction of repeat elements in the assembly. We excluded low complexity regions and small RNA from the analysis (settings –nolow and -norna). REPdenovo v1.0 was run on all pre-processed genomic short reads in default mode with a minimum repeat frequency of 400x (*i.e.* the squared mean genome coverage of 20x; see Results and Discussion section). We then used Bowtie v2 (Langmead and Salzberg 2012) in the sensitive local alignment mode to map all reads to the resulting contigs. The mapping results were then used to calculate mean per-contig coverage with bamtools (Barnett *et al.* 2011). In order to estimate the fraction of the overall genome that corresponds to these highly abundant contigs, we divided the mean contig coverage by the respective contig length and the mean genome-wide coverage. The sum of the resulting normalized bp counts divided by the estimated genome size was then used as an estimate of the fraction of the genome containing highly abundant sequences.

### Genome structure

To analyze genome structure, we compared the genome wide gene synteny of *E. dilemma* and the honey bee. We used the genomic locations of homologous genes (as determined above) of the honey bee and *E. dilemma* scaffolds of at least 100kb length to build haplotype blocks with a minimum length of 1kb. Haplotype blocks included the entire gene span as well as intergenic regions whenever two or more adjacent genes were homologous in both species. We discarded gene annotations from downstream analysis that were recovered as homologous to multiple genomic locations in either species. Furthermore, we excluded *E. dilemma* genes that were recovered as homologous to honey bee scaffolds belonging to unknown linkage groups.

### Data availability

The *E. dilemma* genome assembly *Edil_v1.0*, the annotation, and the original gene set *Edil_OGS_v1.0* are available for download via NCBI [Bioproject: PRJNA388474], Beebase (Elsik *et al.* 2016), the i5k NAL workspace [https://i5k.nal.usda.gov/euglossa-dilemma] (i5K Consortium 2013), and the Ramirez Lab website. The raw reads are available via NCBI [Bioproject: PRJNA388474]. The published raw transcriptome sequence reads are available at the NCBI Sequence Read Archive [SRA: SRX765918] (Brand *et al.* 2015).

## Results and Discussion

### Whole-genome assembly

The *E. dilemma* genome assembly resulted in 22,698 scaffolds with an N50 scaffold length of 144Kb and a total length of 588Mb (Table 1). This represents 18% of the k-mer based estimated genome size of 3.2Gb. Of all sequence reads, 68% aligned to the genome assembly, of which 56% aligned more than once. Further, the k-mer frequency spectrum based on all sequencing reads was strongly positively skewed indicating the presence of highly repetitive sequences in the read set (Figure 1b). Based on the k-mer frequency spectrum, 87.7% of the genome was estimated to be repetitive. This suggests that the genome of *E. dilemma* consists largely of highly repetitive sequences, explaining the low consecutive assembly length, the high assembly fragmentation, and the high fraction of sequence reads mapping multiple times to the assembly. The mean per-base coverage was estimated to be comparatively low in comparison to previous bee genome assemblies, with 19.7x based on the pre-processed reads and estimated genome size (Kocher *et al.* 2013 95.65x coverage; Kapheim *et al.* 2015 120x - 272.3x coverage). Total genomic GC content was 39.9%, and thus similar to previously sequenced bee genomes ranging between 32.7% and 41.5% (Table 1) (Kocher *et al.* 2013; Elsik *et al.* 2014; Kapheim *et al.* 2015).

**Table 1.** *E. dilemma* genome assembly statistics in comparison to previously published bee genomes.

| Species | N50 | N25 | Longest scaffold | Scaffolds | Assembly length | %GC | Predicted genes | Ref. |
| --- | --- | --- | --- | --- | --- | --- | --- | --- |
| *Euglossa dilemma* | 143,590 | 1,417,006 | 10,108,120 | 22,698 | 588,199,720 | 39.94 | 15,904 | 1 |
| *Eufriesea mexicana* | 2,427 | 443,231 | 4,677,300 | 3,522,543 | 1,031,837,970 | 41.38 | 12,022 | 2 |
| *Apis mellifera* | 997,192 | 1,922,192 | 4,736,299 | 5,644 | 234,070,657 | 32.70 | 15,314 | 3 |
| *Melipona quadrifasciata* | 68,085 | 1,896,322 | 12,087,087 | 38,604 | 507,114,161 | 38.88 | 14,257 | 2 |
| *Bombus impatiens* | 1,399,493 | 2,389,513 | 5,466,090 | 5,559 | 249,185,056 | 37.75 | 15,896 | 4 |
| *Lasioglossum albipes* | 616,426 | 1,130,413 | 3,533,895 | 41,433 | 341,616,641 | 41.50 | 13,448 | 5 |

N50 and N25 indicate the length of the shortest scaffold of those including 50% and 25% of the base pairs in a genome assembly. References (Ref.): 1: This study, 2: Kapheim et al. 2015, 3: Elsik et al. 2014, 4: Sadd et al. 2015, 5: Kocher et al. 2012.

Despite the fragmentation of the genome assembly representing less than 20% of the estimated genome size, CEGMA analysis revealed complete assemblies of 231 out of 248 core eukaryotic genes (93.2% completeness). Similarly, BUSCO analysis revealed that 1007 out of 1066 highly conserved arthropod genes were completely assembled (94.4% completeness). The BUSCO analysis detected the duplication of a fraction of 4.8% of the benchmark single-copy ortholog genes in the genome, which is similar to the fraction of arthropod single-copy orthologs found to be duplicated in the honey bee genome (6.9%, Weinstock *et al.* 2006).

Our gene prediction approach generated a comprehensive official gene set including 15,904 protein-coding genes (Table 1). Of these gene models, 11,139 were derived from homology-based predictions, representing 73% of the 15,314 honey bee genes used for annotation. These annotations are well within or exceeding previous bee genome assemblies, and are similar to those reported for the other orchid bee genome available (Table 1) (Kocher *et al.* 2013; Elsik *et al.* 2014; Park *et al.* 2015; Sadd *et al.* 2015; Kapheim *et al.* 2015).

The CEGMA and BUSCO analysis and the gene annotation results suggest that the gene-coding fraction of the *E. dilemma* genome was properly assembled, despite the large estimated genome size and comparatively low per-base sequencing coverage. However, genetic material obtained from natural populations as in our study can lead to the fragmentation of assemblies due to high nucleotide diversity (Kajitani *et al.* 2014). In addition, high genetic diversity in the underlying genetic material can lead to a high number of false duplicates due to multiple incorporation of divergent genomic regions in genome assemblies (Kelley and Salzberg 2010). Our BUSCO analysis suggests that the assembly did not produce an unusually high fraction of duplicated benchmark single-copy orthologs, indicating a relatively low abundance of false duplicates. The observed fragmentation in our assembly is thus likely to be primarily the result of repetitive genomic elements, and less likely the result of low coverage or high nucleotide diversity in the genetic material used for sequencing.

Overall, our results suggest that our approach was sufficient to produce a high quality official gene set. The homology-based approach we used resulted in the majority of annotated genes in the official gene set with a known homology to honey bee genes (Table S1). This genomic resource will facilitate genome-wide expression studies including gene ontology analyses and comparative gene-set analyses among insects.

### Mitochondrial Genome assembly

The recently published transcriptome assembly used for the reconstruction of the mitochondrial genome contained four scaffolds between 1,222bp and 4,188bp long with a total consecutive length of 11,128bp (Figure 2). This corresponds to about 75% of the estimated length of the mitochondrial genome, based on other corbiculate bee species (Crozier and Crozier 1993; Cha *et al.* 2007). The *E. dilemma* mitogenome fragments contained 5 out of 22 tRNAs, 11 out of 13 protein coding genes of which two were only partially recovered, and the 16S rRNA gene. Within scaffolds all genes showed the known hymenopteran gene order and orientation, while the orientation of the 5 tRNAs detected was identical to those in the honey bee (Crozier and Crozier 1993; Cha *et al.* 2007). Attempts to complete the mitochondrial genome using the nuclear genome assembly yielded no improvement of the assembly (data not shown). Accordingly, the mitochondrial genome is entirely derived from transcriptomic sequences.

**Figure 2.**
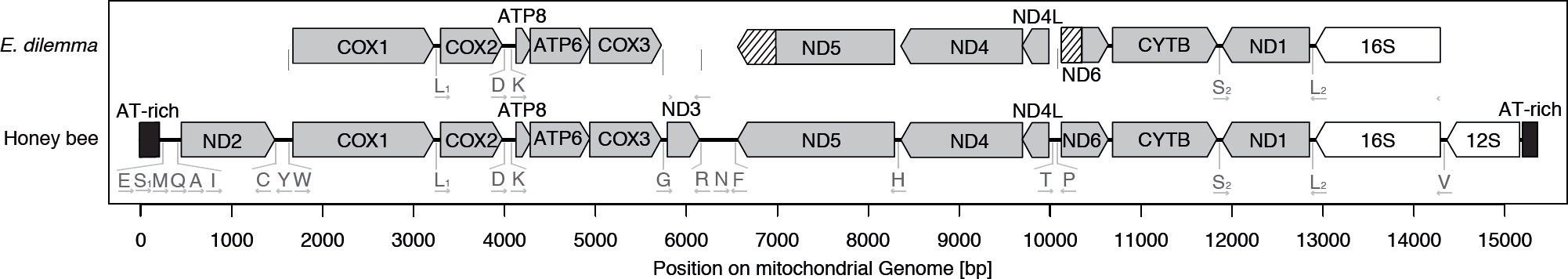
Mitochondrial genome reconstruction. The structure of the honey bee mitochondrial genome and information of the homologous reconstructed parts of the *E. dilemma* mitochondrial genome. Non-reconstructed parts of incompletely reconstructed genes are hatched.

The high success in mitochondrial gene reconstruction is likely due to the nature of the analyzed transcriptome data. Short intergenic regions as well as polycistronic mitochondrial mRNA likely lead to the assembly of multiple genes into single scaffolds. The A-T rich region is completely missing as well as the ND2 and 12S rRNA genes flanking the region in insect mitogenomes. This unrecovered region also contains a high number of tRNAs in the honey bee, which could explain the low number of recovered tRNAs in *E. dilemma*. While the partial mitochondrial genome assembly is only 75% complete, it represents the first mitogenome for the group of orchid bees and will thus be a valuable resource for future phylogenetic analyses within the lineage and between more distantly related bee taxa.

### Repetitive elements

#### Tandem Repeats

We detected 76,001 microsatellite loci with a consecutive length of 2,291,067 bp. Minisatellites with motif lengths from 7bp to 1000bp were less numerous in the genome (67,323 loci), totaling 13,343,515bp. Accordingly, tandem repeats represent 3.86% of the genome assembly, suggesting that they contribute only a small proportion to the overall genome size (Figure 1c).

#### NUMTs

We detected fragments with similarity to the draft mitochondrial genome on 129 scaffolds totaling a length of 150,670 bp. The fragments had a mean length of 764.8 bp and a mean similarity of 91.5% to the mitogenome. This suggests that these fragments are not derived from the mitochondrial genome and represent actual NUMTs. A total of 39 scaffolds carried multiple fragments with high similarity to the mitogenome with a concatenated length of up to 6566bp, suggesting that respective NUMTs might have originated from larger fragments of the mitogenome. In total, only 0.04% of the whole-genome assembly had hits to the mitogenome (Figure 1c). This is likely an underestimate, due to the incompleteness of the reconstructed mitochondrial genome. Nevertheless, NUMTs likely represent only a small fraction of the whole nuclear *E. dilemma* genome. Previous analyses have shown a high density of NUMTs in the honey bee in comparison to other insect genomes totaling about 0.1% of the overall genome size (Pamilo *et al.* 2007). Accordingly, given the NUMT content detected in *E. dilemma*, it is possible that a comparatively high NUMT density is a common feature of corbiculate bee genomes.

#### TEs

The RepeatModeler analysis revealed a total of 566 repeat element families in the genome assembly of which 142 (25.1%) belonged to known TE families including 106 DNA transposons and 36 retroelements, while the remaining 424 (74.9%) repeat element families could not be classified into known TE families (Table2; Supplemental Table 2). Using the 566 newly detected repeat element families as the input database for RepeatMasker, we annotated a total of 597,369 elements in the genome assembly of which 74,513 (12.5 %) were derived from the 142 classified TE families. The remaining 522,856 (87.5 %) elements belonged to the unclassified repeat element families. In total, all annotated repeat elements had a cumulative length of 163,384,833 bp corresponding to 38.48% of the total genome assembly. The majority was derived from unclassified (*i.e.* unknown) repeat element families corresponding to a total of 32.6% of the genome assembly (Figure 1c). Similarly, the read-based *de novo* repeat analysis in REPdenovo revealed 27,636 contigs derived from k-mers with a minimum coverage of 400x (*i.e.* the squared mean genome-wide coverage), which includes 831,433,228 bp (26%) of the estimated 3.3 giga base genome.

**Table 2.** Transposable element repeat class analysis.

| Repeat element family | Number unique elements | Total number elements in assembly | Cumulative length [bp] | Percent of genome assembly<sup>†</sup> |
| --- | --- | --- | --- | --- |
| **Class I - Retrotransposons** | 36 | 12989 | 8843560 | 2.1 |
| **Class II - DNA Transposons** | 106 | 61524 | 16133727 | 3.8 |
| **Total Classified Transposons** | 142 | 74513 | 24977287 | 5.9 |
| **Unclassified** | 424 | 522856 | 138407546 | 32.6 |
| **Total Repeat Families** | 566 | 597369 | 163384833 | 38.5 |

<sup>†</sup>: The percentage was calculated excluding all stretches of N in the scaffolds

The detected high fraction of the genome associated with repetitive element families in *E. dilemma* is not surprising given that large genome sizes are often associated with elevated TE activity and TE content. Similar patterns have been observed in the genomes of diverse lineages, ranging from unicellular eukaryotes to complex multicellular organisms like plants, invertebrates and vertebrates (Kidwell 2002). However, TEs are fast evolving and highly specific to their host lineages, which leads to large underestimates of genomic TE content in previously unstudied lineages (Chalopin *et al.* 2015; Platt *et al.* 2016). This likely explains the large fraction of unclassified repetitive element families that we detected in our genome assembly. Further, the remaining high fraction of unknown genome content (49.15%, Figure 1c) may have resulted from undetected repetitive elements with no similarity to elements from other genomes sequenced previously. The only publicly accessible bee repeat element families are derived from the honey bee, a species with a comparatively small genome (0.23Gb) and low TE diversity and content (Weinstock *et al.* 2006; Kapheim *et al.* 2015). Our efforts to annotate TEs based on known honey bee elements did not improve the TE annotation (data not shown). Overall, our analysis suggests that a large proportion of the *E. dilemma* genome is repetitive. This result is similar to the results obtained in the orchid bee *Eufriesea mexicana*, which has an estimated repetitive genome content of 31% (Kapheim *et al.* 2015).

#### Genome structure

Of the 22,698 *E. dilemma* scaffolds, 580 were at least 100kb in length and used for synteny analysis with the honey bee genome. A total of 356 of these scaffolds carried at least one gene annotation with known homology to the honey bee, and 329 of these *E. dilemma* scaffolds were homologous to honey bee scaffolds with known linkage group (LG) association (Table S1). Of these scaffolds, 272 (83%) showed ≥95% syntenic homology (Figure 1d). Overall, the detected syntenic linkage blocks cover 222MB of scaffold length with homology to the honey bee representing 85% of the 329 filtered scaffolds. This suggests that the genomic architecture is very similar between *E. dilemma* and the honey bee, representing a high level of conservation during the 80 million years since the two lineages diverged. Further, our results support a recent comparative analysis of the honey bee and the bumblebee genomes, which revealed high conservation of genomic synteny (Stolle *et al.* 2011). In comparison, in previous studies gene synteny was found to be less conserved in other insect groups. Extensive local shuffling of gene order beginning on the time scale of 20 - 40 million years evolutionary distance was described for dipterans, moths, and ants resulting in 60-70% genome wide synteny after approximately 60 million years of divergence time in flies and ants (Clark *et al.* 2007; d′Alençon *et al.* 2010; Obbard *et al.* 2012; Simola *et al.* 2013; Neafsey *et al.* 2015; Nygaard *et al.* 2016). Together, these results support a general pattern of slow evolution of gene synteny in corbiculate bees, independent of the fraction of repetitive genome content.

## Conclusion

The genome assembly of the orchid bee *E. dilemma* that we present here is of high quality, despite its large genome size (estimated to be 3.3Gb). The 15,904 gene annotations provide a comprehensive set of genes with known homology to the honey bee, facilitating future gene ontology and functional genomic analyses. While we were unable to annotate the mostly repetitive majority of the genome assembly with our approach, the provided sequence reads will be useful for future analyses of repetitive genetic elements in the genome. The nuclear and mitochondrial draft genomes represent a valuable genomic resource for the community of bee geneticists. This genomic resource will likely prove valuable in genetic and functional genomic analyses dealing with the ecology, evolution, and conservation of orchid bees. Furthermore, the genome of the facultatively eusocial *E. dilemma* will be helpful in the study of the evolution of eusociality, due to its taxonomic placement as the sister lineage to the three obligately eusocial corbiculate bee tribes including stingless bees, bumblebees, and honey bees.

## Acknowledgements

We would like to thank Yannick Wurm and one anonymous reviewer for comments that improved the manuscript. We would like to thank Gene E. Robinson, Guojie Zhang, and the BGI for financial support for genome sequencing. P.B. was supported by a fellowship of the German academic exchange service (Deutscher Akademischer Austauschdienst, DAAD) for parts of the project. SRR received support from the National Science Foundation (DEB- 1457753). This work used the Vincent J. Coates Genomics Sequencing Laboratory at UC Berkeley, supported by NIH S10 Instrumentation Grants S10RR029668, S10RR027303, and OD018174.

